# Transcranial alternating current stimulation at theta frequency to the parietal cortex impairs associative, but not perceptual, memory encoding

**DOI:** 10.1101/2020.11.19.387407

**Authors:** Alyssa Meng, Max Kaiser, Tom de Graaf, Felix Duecker, Alexander T. Sack, Peter de Weerd, Vincent van de Ven

## Abstract

Neural oscillations in the theta range (4-6 Hz) are thought to underlie associative memory function in the hippocampal-cortical network. While there is ample evidence supporting a role of theta oscillations in animal and human memory, most evidence is correlational. Non-invasive brain stimulation (NIBS) can be employed to modulate cortical oscillatory activity to influence brain activity, and possibly modulate deeper brain regions, such as hippocampus, through strong and reliable cortico-hippocampal functional connections. We applied high-definition transcranial alternating current stimulation (HD-tACS) at 6 Hz over left parietal cortex to modulate brain activity in the putative cortico-hippocampal network to influence associative memory encoding. After encoding and brain stimulation, participants completed an associative memory and a perceptual recognition task. Results showed that theta tACS significantly decreased associative memory performance but did not affect perceptual memory performance. These results show that parietal theta tACS modulates associative processing separately from perceptual processing, and further substantiate the hypothesis that theta oscillations are implicated in the cortico-hippocampal network and associative encoding.

## Introduction

Associative memory is commonly defined as a memory system that constructs relationships between initially unrelated items (Suzuki, 2008; Tulving, 1972). The ability to form associations is core to episodic memory function (Kahana, Howard, & Polyn, 2008; Lang, Gan, Alrazi, & Monchi, 2019), and allows us to remember details of specific events and recall abstract information, required for daily tasks and interactions (Herweg, Solomon, & Kahana, 2020). The study of neural mechanisms underlying associative memory has traditionally focused on hippocampus and its central role in widespread brain networks. Given the rapid and flexible nature of associative memory, a dynamic mechanism for neural communication is required. In this regard, neural oscillatory rhythms are of particular interest because they are thought to contribute to the communication between neuronal populations that are simultaneously engaged in cognitive processes (Buzsáki & Draguhn, 2004; X.-J. Wang, 2010). In particular, oscillatory activity in the theta range (4 – 8 Hz) has been extensively researched in animal and human cognition (Buzsáki, 2002; Jacobs, 2014; Meador et al., 1991), as it has been hypothesized to play an integral part in hippocampal function during spatial navigation and associative memory processing (Buzsáki, 2002; Skelin, Kilianski, & McNaughton, 2019; Wallenstein, Hasselmo, & Eichenbaum, 1998). The success of memory encoding seems to rely on the increase of theta power in hippocampal regions (Burke et al., 2013; Khader, Jost, Ranganath, & Rösler, 2010; Sato & Yamaguchi, 2007) although decreased theta power for subsequently successfully remembered items has also been observed (Crespo-García et al., 2016; Fellner et al., 2016; Lega, Jacobs, & Kahana, 2012).

The study of hippocampal oscillatory mechanisms in associative memory, particularly in humans, can be challenging, as its location deep within the brain precludes direct noninvasive measurement of neural activity. Moreover, measuring oscillations does not always allow assessment of causal contributions, which ideally involves direct experimental manipulation of the neural mechanisms of interest. Such manipulation can be achieved with non-invasive brain stimulation (NIBS), by taking advantage of local (i.e., cortical) and network functional connectivity in order to investigate cortical and subcortical memory effects. Recent transcranial magnetic stimulation (TMS) studies showed that cortical stimulation may alter activity and connectivity within a hippocampal-cortical network, which could in turn lead to enhanced memory performance (Hermiller, Chen, Parrish, & Voss, 2020a; Hermiller, VanHaerents, Raij, & Voss, 2019; J. X. Wang et al., 2014). Here, experimentally induced changes in cortical activity may spread to more remote areas through default functional connectivity (Kim, Ekstrom, & Tandon, 2016). Particularly, the lateral parietal cortices have been shown to be strongly and reliably functionally connected to the hippocampus (Buckner, Andrews-Hanna, & Schacter, 2008; van de Ven, Wingen, Kuypers, Ramaekers, & Formisano, 2013; Vincent et al., 2006). The ease of stimulation of parietal cortex, combined with the limited discomfort from transcranial stimulation over these areas, further exemplifies how this area can serve as an important node for network-based stimulation in associative memory.

A different NIBS technique, transcranial alternating current stimulation (tACS) possesses the ability to modulate neural networks through a pre-determined exogenous frequency (Helfrich et al., 2014; Herrmann, Rach, Neuling, & Strüber, 2013; Reed & Kadosh, 2018). To date, tACS has been used to study the oscillatory mechanisms in associative memory in only a limited number of studies. In one study (Lang et al., 2019), participants encoded the associations between arbitrary face-scene pairs while undergoing theta-tACS (or anodal transcranial direct current stimulation (tDCS) or sham stimulation) over the fusiform area, after which they completed an associative memory test. Results showed that theta tACS during encoding enhanced associative memory performance, compared to tDCS or sham. However, tACS also improved correct rejections of lure items, which suggests a more general cognitive enhancement effect. Previous studies reported theta oscillatory activity in human and animal prefrontal and sensory cortex during working memory procedures (Lee et al., 2005; Raghavachari et al., 2006; Sauseng, Griesmayr, Freunberger, & Klimesch, 2010), as well as during perceptual processing (Helfrich et al., 2018; Landau, Schreyer, Van Pelt, & Fries, 2015; Somer, Allen, Brooks, Buttrill, & Javadi, 2020), which support the contribution of various cognitive processes to associative memory performance. Indeed, a number of studies reported enhanced working memory performance after theta tACS was administered over parietal (Jaušovec & Jaušovec, 2014) or prefrontal cortex (Alekseichuk, Turi, de Lara, Antal, & Paulus, 2016), while tACS-induced frontoparietal inter-regional desynchronization at 6 Hz resulted in decreased working memory performance (Alekseichuk, Pabel, Antal, & Paulus, 2017). However, another study in which theta tACS was administered over left prefrontal cortex during associative memory encoding did not alter memory performance in young healthy participants (Klink, Peter, Wyss, & Klöppel, 2020).

In sum, findings of non-invasive theta stimulation on associative memory remain inconclusive. A number of factors need to be addressed to further our understanding. First, the effect of theta stimulation on associative memory may be strengthened when applied to cortical areas that are strongly connected to the hippocampal-cortical networks. Based on recent TMS studies, the parietal cortex may be a good candidate for tACS administration. However, a direct application of theta tACS over parietal cortex in order to selectively alter associative memory processing has not yet been reported. Second, the specificity of theta stimulation on associative memory requires that non-associative cognitive functions were not modulated. This is especially relevant given that parietal cortex plays a role in associative memory as well as perceptual processing (Cabeza et al., 2011; Culham & Kanwisher, 2001; Gottlieb, 2007). Parietal theta tACS could thus modulate associative processing indirectly by modulating sensory or attentional encoding, which in turn could modulate associative and non-associative memory performance, such as perceptual recognition. Supporting the functional specificity of theta oscillatory activity in associative memory would require a theta tACS effect on associative memory in the absence of an effect on perceptual memory.

In the current study, we administered high-definition tACS at 6 Hz over left parietal cortex in a within-subject sham-controlled design while participants encoded a face-scene pairing task. After the encoding phase and tACS, participants completed two memory tests that measured knowledge about the item pairings (i.e., associative memory) and item identity (i.e., perceptual memory). We hypothesized that parietal theta tACS, compared to sham stimulation, would affect associative, but not perceptual memory performance.

## Methods

### Participants

We recruited twenty healthy volunteers (mean age [SD] = 21.68 years [2.84], 12 females) from the academic environment of Maastricht University via an online recruitment system. All participants were pre-screened using a non-invasive brain stimulation screening form for eligibility, health risks, and adverse effects. Potential participants who reported any NIBS contraindications during their screening phase were excluded. All participants gave their written informed consent prior to participation. The study was approved by the ethics review committee of psychology and neuroscience (ERCPN) of the Faculty of Psychology and Neuroscience (FPN) at Maastricht University. Participant safety and comfort were monitored throughout brain stimulation in order to catch any adverse effects due to stimulation. Participants were compensated with a gift voucher or course credit.

### Task Procedure

Participants completed a visual associative memory task, the “Face and Scene Task” (FAST; (Lang et al., 2019), which was programmed in Psychopy (Peirce et al., 2019). The task consisted of an encoding and a test phase. During the encoding phase of the task, participants were presented with image pairs that comprised a neutral face and a natural scene. Neutral faces of all ethnicities and ages were gathered from the Chicago Face Database (CFD Version 2.0.3; (Ma, Correll, & Wittenbrink, 2015)). The faces did not have easily distinguishable features such as piercings or tattoos. All scene images were taken from an online image database (www.pixabay.com).

On each trial, participants saw a pair of images consisting of a neutral face presented on the left side of the screen, and a scene presented on the right side of the screen. They were given 8 seconds to indicate how the person in the presented image would appraise the corresponding scene. Participants were instructed to provide their judgement via a rating scale that was visualised as a scaled slider (range −3 to 3) below the image pair to ensure the participants were actively encoding (Figure 1). The image disappeared after a button press. Each face-scene pair was followed by a 100 ms inter-stimulus interval during which a fixation cross was presented. The encoding phase consisted of 45 face-scene pairs in total, which were separated into 5 blocks of 9 image pairs. Each block was separated by a 5 second interval.

**Figure 1.**
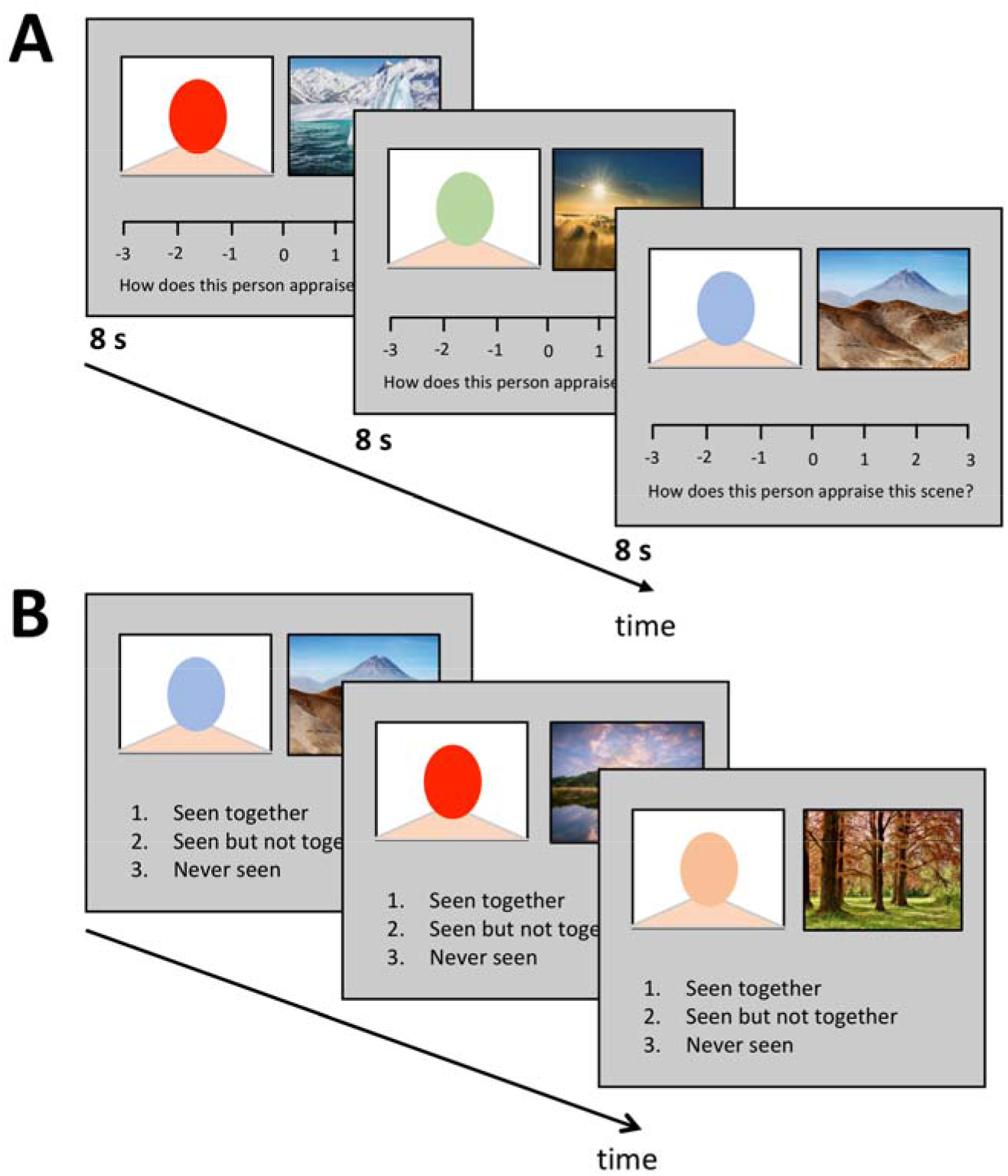
Encoding and retrieval phase. During the encoding phase (A), participants saw face-scene pairs and indicated how positively or negatively they thought that the person appraised the scene. During the memory test phase (B), participants saw face-scene pairs and had to indicate whether the items were previously seen as a pair (“1: seen together”), seen as items of different pairs (“2: seen but not together”), or were not seen in any way before (“3: never seen”). Participants were given 8 seconds to encode and rate each face-scene pair. In the retrieval phase, participants were given an unlimited time to respond.

Each participant performed two versions of this task, which were identical in design but used non-overlapping sets of face and scene images. Each encoding task was presented during either 6 Hz or sham tACS stimulation. In order to familiarise the participants with the task, an abridged version of 15 novel pairs were given as training prior to the experimental conditions and standardised instructions were presented at the beginning of each version.

Following the encoding task, participants completed a video game as a distraction task for 5 minutes (www.tetris.com) in order to diminish the recency effect of short-term memory and minimise possible stimulation after-effects. The task is believed to be engaging enough to prevent effortful rehearsal.

After the distraction task, participants completed the memory testing phase, which comprised an associative and a perceptual recognition task. In the associative recognition task, participants saw 45 presentations of either 1) the original face-scene pairs seen during encoding, 2) pairs of faces and scenes that were previously seen but taken from different encoding pairs, or 3) pairs of novel images that were not previously seen during encoding. Participants were asked to indicate whether they had “seen together”, “seen but not together”, or “never seen” with the keys ‘1’, ‘2’, and ‘3’ respectively (Figure 1). Participants were asked to make keyboard responses with their dominant hand. There was no time limit.

Associative retrieval blocks were intermixed with the perceptual recognition task, in which a series of novel or previously seen images were presented for 200 ms, after which participants were asked if they recognised the images (Yes/No responses). Per block, participants answered 9 associative retrieval image pairs and then answered 9 perceptual recognition images (Figure 2). These blocks would alternate in an A-B-A-B format.

**Figure 2.**
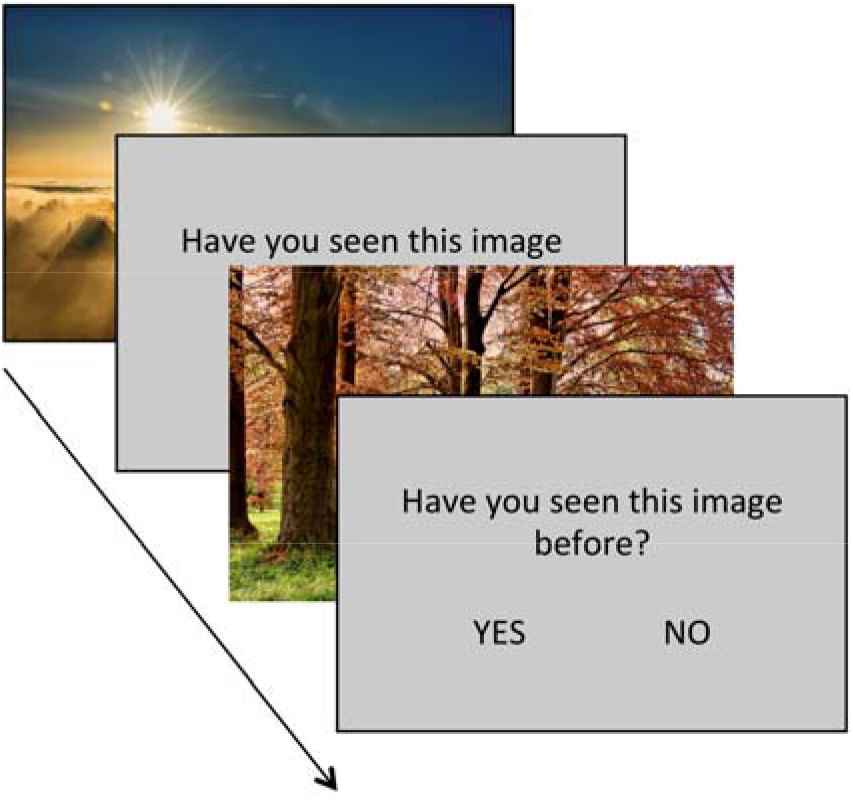
Perceptual recognition task. Participants judged (“yes” or “no”) whether single items were previously seen during the FAST encoding phase (see Figure 1). Each image was presented for 200ms, followed by a 100ms interstimulus. Participants were given an unlimited time to answer whether they had seen the image before (yes/no).

The order of the tACS conditions was counterbalanced across participants. Participants rested their head in a fixed chin rest before testing began. The experiment lasted approximately 90 minutes per participant, which included the time required to attach the electrodes. Each participant underwent the sham and theta stimulation condition on the same day, with approximately 30 min between the two stimulation periods.

### tACS parameters

tACS was applied using a neuroConn DC-STIMULATOR TES device, using a concentric stimulation setup for high-definition focal stimulation, which included an inner circular electrode (3 cm diameter) placed inside an outer electrode (11 cm diameter). EEG caps were used to locate the area of stimulation, P3, for each individual, as this site has been shown to have extensive connections with the hippocampal network (Freedberg et al., 2019; Hermiller et al., 2019; J. X. Wang et al., 2014). Each participant was examined for any microlesions on the scalp before electrode attachment. Conductive paste (Ten20) was used to optimise conductivity between the electrodes and scalp to below 10kΩ for each participant.

In the theta condition, the frequency of stimulation was set to 6 Hz for 5400 cycles (15 minutes x 60 seconds x 6 Hz) at an intensity of 2 mA (peak-to-peak). Participants were familiarised with the sensation of tACS in a short stimulation session to determine if the stimulation was tolerable. One participant indicated discomfort, and stimulation intensity was reduced for this participant to 1.8 mA. The ramp-up and ramp-down durations were set to 30 seconds in the theta condition. In the sham condition, stimulation lasted for 300 cycles (50 seconds) before ramping down in order to create an initially indistinguishable state to mask the conditions.

Due to the difference in continuous somatosensory sensations between the two tACS conditions, participants could theoretically discern the theta stimulation from the sham stimulation. To verify whether this was the case, participants were asked to verbally identify the differences in stimulation conditions post-hoc at the end of the experiment. None of the participants informed us they were able to confidently distinguish the theta stimulation from the sham.

### Data Analysis

Statistical analysis was performed in JASP (Team, 2020). Of the twenty participants, data of one participant was excluded for having incomplete datafiles as a result of an error in the experiment program code. Data of another participant were excluded after reporting suffering from fatigue before and during the experiment. Of the remaining datasets, trials with a clear delayed response (> 2 x SD) were excluded from the analysis.

We analysed memory performance by calculating memory sensitivity (d’;(Macmillan & Creelman, 2005; Stanislaw & Todorov, 1999) using hit rates and false alarm rates. For the associative memory test, hit rate was calculated as the proportion of face-scene pairs shown during the encoding task, which were correctly identified as “seen as a pair”. The false alarm rate was identified as the proportion of face-scene pairs that were wrongly judged to have been seen as a pair during the encoding task. For the perceptual memory test, hit rate was calculated as the proportion of previously shown single items that were correctly judged as seen before, and false alarm rates as the proportion of novel items that were erroneously judged as seen before. Statistical comparisons between tACS conditions were carried out using paired sample *t*-tests, and repeated measures ANOVAs were used to test whether the within-subject effects correlated with covariates Age, Sex and Order of tACS condition. Effects were deemed significant at a p-value of 0.05.

As our hypotheses included null effects, we calculated Bayes Factor to quantify the strength of the evidence in favour of no effect due to tACS, that is, BF_10_ < 1 (Wagenmakers, 2007). Bayes Factor calculation was performed using JASP.

## Results

### Theta-based tACS effects on associative memory responses

Associative d’ memory performance during theta-tACS (mean [SE] = 1.32 [0.13]) was significantly lower than associative d’ during sham-tACS (1.61 [0.16]; *t*(17) = −2.517, *p* = .022, Cohen’s *d* = −0.59; see Figure 3). A repeated-measures ANOVA with tACS (Theta, Sham) as within-subject factor showed that this effect did not interact with between-subject covariates Age (P = 0.23) or Sex (P = 0.36). We also tested for carry-over effects of theta stimulation on the encoding during the subsequent sham stimulation session by using tACS Order (theta followed by sham vs. sham followed by theta) as between-subject covariate, which did not significantly interact with the within-subject tACS effect on associative memory (P = 0.64).

**Figure 3.**
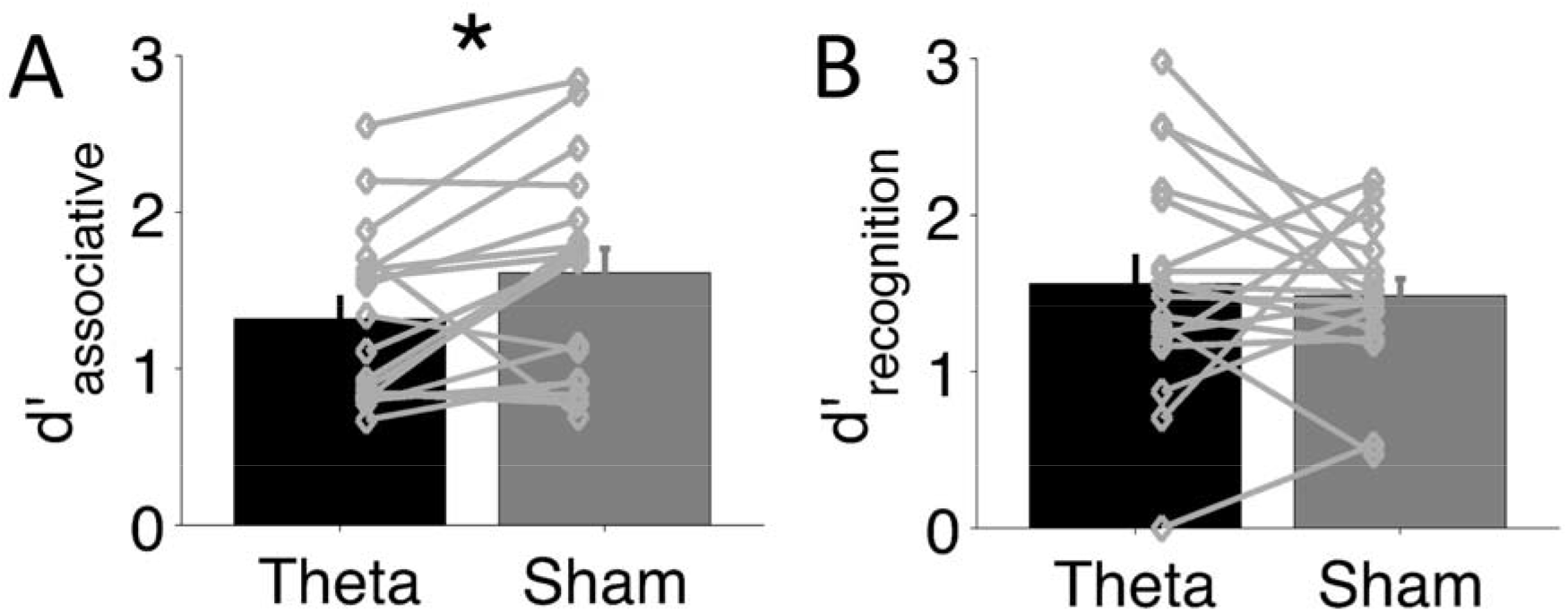
Results. Bar-plots show mean d’ for the Theta (black bars) and Sham (grey bars) tACS conditions for the associative memory (A) and perceptual recognition tasks (B). Grey dots and lines indicate paired values of individual participants. Error bars depict standard error of the mean (SEM), * p < 0.05.

To further investigate the effect of tACS on d’, we conducted post-hoc comparisons for hit rate and false alarm rate between the two tACS conditions. The hit rate was significantly higher in the sham condition (67.0% [0.05]) compared to the theta condition (53.7% [0.05]) for associative memory (*t*(17) = −3.286, *p* = 0.004, Cohen’s d = −0.78). False alarm rates for associative memory performance revealed no significant difference between the sham (16.2% [0.02]) or theta condition (12.1% [0.02]: *t*(17) = −1.428, *p* = 0.171). These findings corroborated that theta-tACS impaired associative memory performance.

### Theta-based tACS effect on perceptual item recognition

Next, we analysed d’ scores for the perceptual recognition test, which was performed on the same visual material as for the associative memory test. We found no significant difference in perceptual item recognition between the theta (1.56 [0.72]) and sham conditions (1.48 [0.47]; *t*(17) = 0.462, *p* = 0.65). As we hypothesized no tACS effect on perceptual recognition, we calculated Bayes Factor (BF) for this comparison. We found that BF_10_ = 0.44, which indicated that the evidence was in favour of no effect.

### Comparison between associative and perceptual memory performances

To analyse whether tACS affected associative and perceptual recognition memory differently, we conducted four further analyses. First, we correlated d’ for associative memory and perceptual recognition performance for the sham condition to determine whether baseline performance in the two tasks were related. This correlation was not significant (r = 0.28, *p* = 0.26). Calculation of Bayes Factor showed that the evidence was in favour of performance of the two tasks not being related (BF_10_ = 0.52). This result confirms that the associative and perceptual memory task rely on different underlying neural mechanisms. Second, we compared d’ of the sham condition of the two memory tasks, using a paired-samples *t*-test, and found that they were not significantly different (*t*(17) = 0.78, *p* = 0.78; BF_10_ = 0.25), which indicates that the tasks were equally difficult. Third, we correlated the performance difference due to tACS (i.e., d’(Theta) – d’(Sham)) between associative and perceptual recognition memory, and found no significant correlation (r = 0.12, *p* = 0.63; BF_10_ = 0.33). Fourth, we tested whether the tACS-induced impairment of associative memory performance was larger than the (null) effect of tACS on recognition memory performance. To model this interaction between tACS and task, we calculated a repeated measures model with tACS and Task as within-subject factors (and Age, Sex and Order as covariates), and found a significant tACS x Task interaction effect (F(1,13) = 7.68, *p* = 0.015, 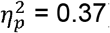). In all, these results indicate that tACS affected associative memory differently from recognition memory performance.

## Discussion

Our results demonstrate that theta tACS over parietal cortex during memory encoding impairs associative memory performance but not perceptual recognition memory of the same task materials. These findings are in line with repeated observations that theta oscillatory activity specifically contributes to associative memory encoding (Burke et al., 2013; Buzsáki, 2002; Herweg et al., 2020; Lega et al., 2012; Wallenstein et al., 1998). The finding that associative memory performance did not correlate with perceptual memory performance further corroborates the notion that the neural correlates of the two memory processes are dissociable. This is of further importance given that the site of parietal stimulation (P3) has been found to play a role in both associative memory and perceptual processing (Cabeza et al., 2011; Culham & Kanwisher, 2001; Gottlieb, 2007). That is, the neural correlates of the two tasks show spatial overlap in parietal cortex. Yet, the dissociable effect of theta tACS on the two memory tasks suggests that theta oscillatory activity may functionally dissociate the two memory processes in parietal cortex. Furthermore, this dissociation cannot be explained by a difference in task difficulty, as performance during the sham conditions of both tasks was similar.

Concurrently, our finding of an associative memory impairment after theta tACS contrasts several previous findings of theta tACS-induced enhanced memory performance. However, comparison between studies is hampered by the heterogeneity in task design and location of brain stimulation. For example, tACS-induced memory enhancement has been reported in working memory studies (Alekseichuk et al., 2016; Khader et al., 2010), but these tasks arguably did not rely on associative processing. Further, the area of stimulation may be of particular importance due to the differential effects of theta power on different neuroanatomical sites (Long, Burke, & Kahana, 2014). For example, the enhanced performance in the face-scene association task after tACS was administered to the fusiform area (Lang et al., 2019) could have resulted from modulation of the encoding of facial features, rather than increased associative memory encoding. While theta oscillatory activity has been observed in these areas in humans and animals (Lee et al., 2005; Raghavachari et al., 2006; Sauseng et al., 2010), they may play a different role in memory processes. In a more recent study, Klink et al. (2020) found no change in associative memory performance in young participants after tACS over the ventrolateral prefrontal cortex, which further exemplifies the inconsistency in results.

Instead, we suggest that our findings are in line with previous studies showing that parietal cortical stimulation modulates neural activity in a hippocampal-cortical network that supports associative memory processing (Hermiller et al., 2020a; Hermiller, Chen, Parrish, & Voss, 2020b; Hermiller et al., 2019; Tambini, Nee, & D’Esposito, 2018; J. X. Wang et al., 2014). More particularly, our findings are in line with the modulation of associative memory, but not recognition memory, after administration of TMS over parietal cortex (Hermiller et al., 2019). We extend these reports by showing that theta tACS can also modulate associative processing within arguably the same hippocampal-cortical network. It is possible that our finding of decreased, rather than enhanced, associative memory may indeed be related to the theta oscillatory pattern of brain stimulation. There is evidence that associative encoding in human individuals is supported by two frequency bands at the extremes of the theta range (3 – 8 Hz), rather than varying around the centre of the theta range (Lega et al., 2012). tACS modulation at 6 Hz could then result in desynchronisation of an individual’s endogenous theta waves, resulting in a disruption during encoding. This postulation is further in line with reports of decreased memory performance following tACS-induced oscillatory phase-desynchronisation (Alekseichuk et al., 2017; Vosskuhl, Strüber, & Herrmann, 2018; Wolinski, Cooper, Sauseng, & Romei, 2018).

The design of our study poses a few limitations to our inferences. We did not include functional brain imaging simultaneous to tACS administration, which prevents us from verifying the effect of parietal tACS on the hippocampal-cortical network. Also, we did not measure EEG simultaneous to tACS administration, which means that we could not empirically test our postulation of theta desynchronization in associative memory. Finally, we did not use a control frequency or location to verify the specificity of theta stimulation of parietal cortex. We are planning to tackle these open issues in future studies.

In conclusion, our tACS study demonstrates that theta oscillations are involved in the encoding of associative memory, while sparing perceptual item recognition and general memory performance. These results extend current knowledge on theta oscillations in the parieto-hippocampal network, and advocate merit in using non-invasive electric brain stimulation to modulate oscillatory activity of associative memory brain networks for current memory research.

## Acknowledgments

We thank Selma Kemmerer with practical assistance with brain stimulation.

## Author contributions

AM, VV: Designed the study; VV: Supervised the study; TdG, FD: Contributed methodology; AM, MK: Collected the data; AM, VV: Analysed the data; AM wrote the first version of the manuscript; all authors contributed to data interpretation and writing/editing of the manuscript.

